# A Goldilocks Principle for the Gut Microbiome: Taxonomic Resolution Matters for Microbiome-Based Classification of Colorectal Cancer

**DOI:** 10.1101/2021.10.22.465538

**Authors:** Courtney R. Armour, Begüm D. Topçuoğlu, Andrea Garretto, Patrick D. Schloss

## Abstract

Colorectal cancer is a common and deadly disease in the United States accounting for over 50,000 deaths in 2020. This progressive disease is highly preventable with early detection and treatment, but many people do not comply with the recommended screening guidelines. The gut microbiome has emerged as a promising target for non-invasive detection of colorectal cancer. Most microbiome-based classification efforts utilize taxonomic abundance data from operational taxonomic units (OTUs) or amplicon sequence variants (ASVs) with the goal of increasing taxonomic resolution. However, it is unknown which taxonomic resolution is optimal for microbiome-based classification of colorectal cancer. To address this question, we used a reproducible machine learning framework to quantify classification performance of models based on data annotated to phylum, class, order, family, genus, OTU, and ASV levels. We found that model performance increased with increasing taxonomic resolution, up to the family level where performance was equal (p > 0.05) among family (mean AUC: 0.689), genus (mean AUC: 0.690), and OTU (mean AUC: 0.693) levels before decreasing at the ASV level (p < 0.05, mean AUC: 0.676). These results demonstrate a trade-off between taxonomic resolution and prediction performance, where coarse taxonomic resolution (e.g. phylum) is not distinct enough but fine resolution (e.g. ASV) is to individualized to accurately classify samples. Similar to the story of Goldilocks and the three bears, mid-range resolution (i.e. family, genus, and OTU) is just right for optimal prediction of colorectal cancer from microbiome data.

**Importance:** Despite being highly preventable, colorectal cancer remains a leading cause of cancer related death in the United States. Low-cost, non-invasive detection methods could greatly improve our ability to identify and treat early stages of disease. The microbiome has shown promise as a resource for detection of colorectal cancer. Research on the gut microbiome tends to focus on improving our ability to profile species and strain level taxonomic resolution. However, we found that finer resolution impedes the ability to predict colorectal cancer based on the gut microbiome. These results highlight the need for consideration of the appropriate taxonomic resolution for microbiome analyses and that finer resolution is not always more informative.

---

Colorectal cancer is one of the most common cancers in men and women and a leading cause of cancer related deaths in the United States (1). Early detection and treatment are essential to increase survival rates, but for reasons such as invasiveness and high screening costs (i.e. colonoscopy), many people do not comply with recommended screening guidelines (2). This prompts a need for low cost, non-invasive detection methods. A growing body of research points to the gut microbiome as a promising target for non-invasive detection of screen relevant neoplasia (SRNs) consisting of advanced adenomas and carcinomas (3, 4). The diagnostic potential of the gut microbiome in detecting SRNs has been explored through machine learning (ML) methods using abundances of operational taxonomic unit (OTU) classifications based on amplicon sequencing of the 16S rRNA gene (3). Recent work has pushed for the use of amplicon sequence variants (ASVs) to replace OTUs for marker-gene analysis because of the improved resolution with ASVs (5). However, it is unclear if OTUs are the optimal taxonomic resolution for classifying SRNs from microbiome data or whether the additional resolution provided by ASVs is useful for ML classification. Topçuoğlu *et al* (6) recently demonstrated how to effectively apply machine learning (ML) methods to microbiome based classification problems and developed a framework for applying ML practices in a more reproducible way. This analysis utilizes the reproducible framework developed by Topçuoğlu *et al* to determine which ML method and taxonomic level produce the best performing classifier for detecting SRNs from microbiome data.

Utilizing publicly available 16S rRNA sequence data from stool of patients with SRNs and healthy controls, we generated taxonomic abundance tables with mothur (7) annotated to phylum, class, order, family, genus, OTU and ASV levels. Using the taxonomic abundance data and the mikropml R package (8), we quantified how reliably samples could be classified as SRN or normal using five machine learning methods including random forest, L2-regularized logistic regression, decision tree, gradient boosted trees (XGBoost), and support vector machine with radial basis kernel (SVM radial). Across the five machine learning methods tested, model performance increased with increasing taxonomic level usually peaking around genus/OTU level before dropping off slightly with ASVs (Supplemental Figure 1). Regardless of the taxonomic level, random forest (RF) models consistently had the largest area under the receiver operating characteristic curve (AUROC). Within the RF model, the highest AUROCs were observed for family (mean AUROC: 0.689), genus (mean AUROC: 0.690), and OTU (mean AUROC: 0.693) level data with no significant difference between the three (p > 0.05, Figure 1A, Supplemental Figure 2). Performance with ASVs (mean AUROC: 0.676) was significantly lower than OTUs (p < 0.01), but comparable to family (p = 0.06) and genus (p = 0.05) levels (Figure 1A). These results suggest that increased resolution improves model performance up to the OTU level where further taxonomic resolution is not necessarily better for identifying individuals with SRNs based on microbiome composition.

**Figure 1:**
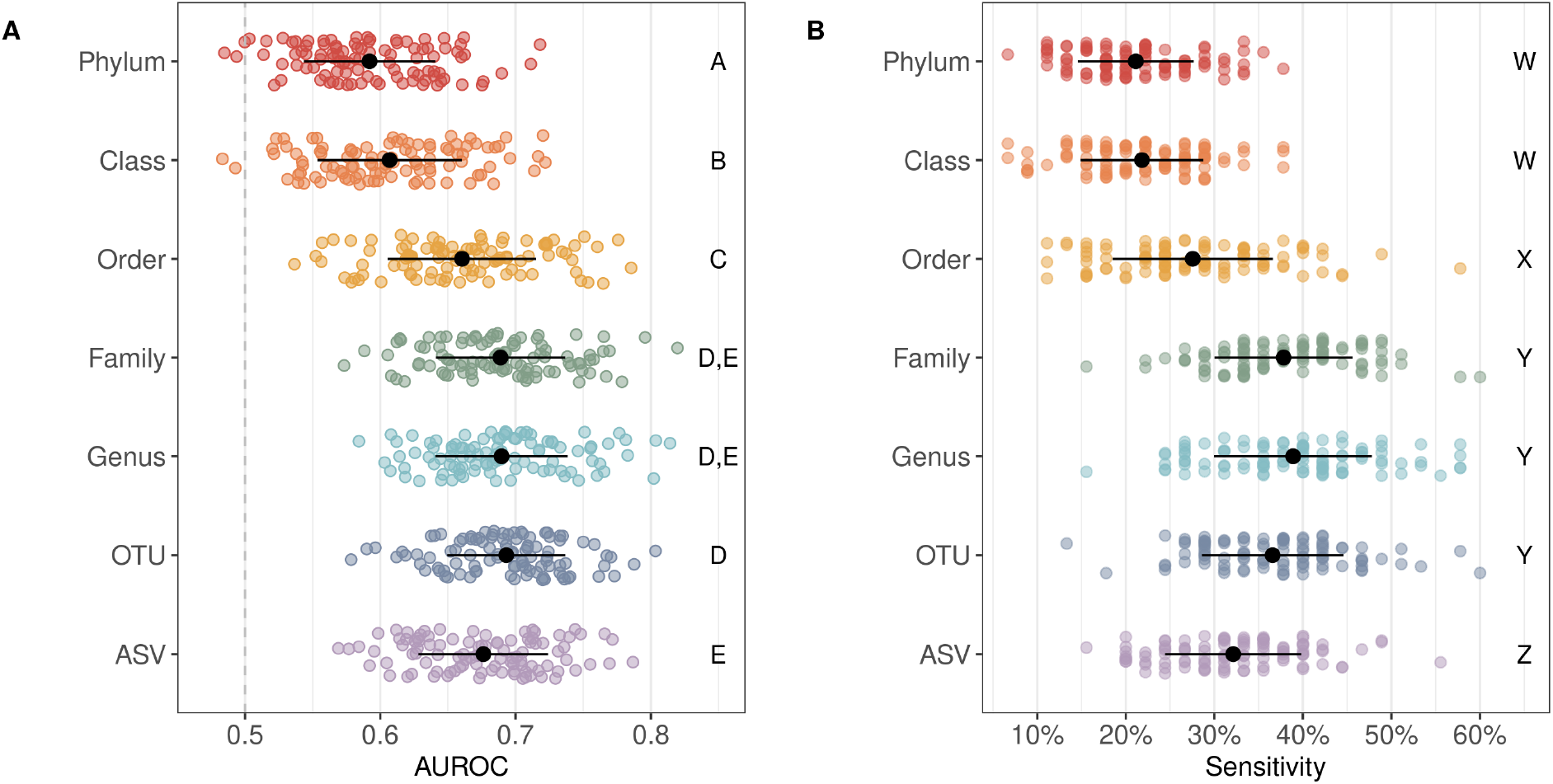
Random Forest Model Performance. **A)** Strip plot of the area under the receiver operating characteristic curve (AUROC) values on the test dataset for 100 seeds predicting SRNs using a random forest model. Black points denote the mean and lines denote the standard deviation. Dashed line denotes AUROC of 0.5 which is equivalent to random classification. Significance between taxonomic levels was quantified by comparing the difference in mean AUROC and is denoted by letters A through E on the right side of the plot; taxonomic levels with the same letter are in the same significance group and are not significantly different from one another. **B)** Strip plot of the sensitivity at a specificity of 90% across the 100 model iterations for each taxonomic level. Black points denote the mean and the lines denote the standard deviation. The letters W through Z on the right side of the plot denote the significance groups.

While comparing AUROC values between models is a useful way to assess the overall model performance, they summarize the performance across all thresholds and can be misleading since models with the same AUROC can have different ROC curve shapes (9). Depending on the intended implementation of the model, one may want to optimize the sensitivity over the specificity or vice versa. In this case, the optimal model will detect as many true positives (people with SRNs) as possible. To further compare the model performance across taxonomic levels we compared the sensitivity of the models at a specificity of 90%. The highest sensitivity values were observed for family (mean sensitivity: 0.38), genus (mean sensitivity: 0.39), and OTU (mean sensitivity: 0.37) level data (p > 0.05, Figure 1B), consistent with the AUROC results. Phylum (mean sensitivity: 0.21), class (mean sensitivity: 0.22), order (mean sensitivity: 0.28), and ASV (mean sensitivity: 0.32) sensitivity values were all significantly lower than family, genus, and OTU sensitivity values (p < 0.05,Figure 1B). This analysis further supports the observation that finer resolution does not improve SRN detection.

One hypothesis for the observation that model performance increases from phylum to OTU level then drops at the ASV level is that at higher taxonomic levels (e.g. phylum) there are too few taxa and too much overlap to reliably differentiate between cases and controls. At the level of genus or OTU there is enough data and variation but at the ASV level, the data is too specific to individuals and does not overlap enough. Examination of the prevalence of taxa in samples at each level supports this idea. A majority of taxa were present in greater than 70% of samples at the phylum (67% of taxa) and class (63% of taxa) levels. The opposite was observed at the OTU and ASV level where 50% and 41% of taxa, respectively, were only present in 20% or less of the samples (Supplemental Figure 3). Of note, the ML pipeline includes a pre-processing step that occurred prior to training and classifying the ML models. For example, perfectly correlated taxa provide the same information to build the model and thus can be collapsed. Additionally, features with zero or near-zero variance across samples were removed. Interestingly, despite starting with 104,106 ASVs, only 478 (0.5%) remained after pre-processing. At the OTU level, 705 of the 20,079 OTUs (3.5%) remained after preprocessing (Table 1). While the resolution provided by ASVs is useful in certain contexts(10, 11), these results suggest that the resolution is too fine for use in machine learning classification of SRNs based on microbiome composition.

**Table 1:** Summary of Features. Overview of the number of features at each taxonomic level before and after preprocessing as described in the methods.

| Taxonomic Level | Number of Features | Number of Features | Percent of Features Kept |
| --- | --- | --- | --- |
|  |  | After Preprocessing | After Preprocessing |
| Phylum | 19 | 9 | 47.4 % |
| Class | 36 | 19 | 52.8 % |
| Order | 65 | 28 | 43.1 % |
| Family | 124 | 54 | 43.5 % |
| Genus | 316 | 115 | 36.4 % |
| OTU | 20,079 | 705 | 3.5 % |
| ASV | 104,106 | 478 | 0.5 % |

A look into the most important taxa at each level for classifying samples revealed some nesting where several genera and their higher taxonomic classifications were in the top 10 most important taxa (Supplemental Figure 4). For example, the genus *Gemella* was an important taxon at the genus and OTU levels and its higher classifications were also important (*Firmicutes* > *Bacilli* > *Bacillales* > *Bacillales Incertae Sedis XI* > *Gemella*). *Fusobacterium* displayed a similar pattern, except that the family level classification (*Fusobacteriaceae*) importance was ranked 16th out of 54 families. In the case of unclassified *Lachnospiraceae*, there were several OTUs with this label that were in the top 10, however at the genus level this taxon was ranked lower in importance (21st out of 115 genera) suggesting there may be some benefit to separating different taxonomic groupings within *Lachnospiraceae*.

These results demonstrate a Goldilocks effect such that consideration of the appropriate taxonomic resolution for utilizing the microbiome as a predictive tool is warranted. In general, we found that finer taxonomic resolution (e.g. ASV) did not add additional sensitivity to predicting SRNs based on microbiome composition. Family, genus, and OTU level data all performed equally. At the ASV level the fine resolution actually impeded model performance due to the sparsity of shared taxa and led to decreased model performance. The tendency for ASV level annotation to split single bacterial genomes into multiple taxa (12) could also be a contributing factor to the sparsity of shared taxa. Additionally, these results indicate that there are not specific individual bacterial strains that are useful to resolve SRNs, rather sets of closely related bacterial taxa. Overall, either family, genus, or OTU level taxonomy appear to perform equally for predicting subjects with SRNs based on the composition of the gut microbiome. A potential benefit of utilizing genus or family level data could be that it may allow for merging data generated from different 16S rRNA gene regions or sequencing platforms.

## Materials and Methods

### Dataset

Raw 16S rRNA gene amplicon sequence data isolated from human gut samples (13) was downloaded from NCBI Sequence Read Archive (SRP062005). This dataset contains stool samples from 490 subjects. Based on the available metadata, samples categorized as normal, high risk normal, or adenoma were labeled “normal” for this analysis and samples categorized as advanced adenoma or carcinoma were labeled as “screen relevant neoplasia” (SRN). This resulted in a total of 261 “normal” samples and 229 “SRN” samples.

### Data processing

Sequence data was processed with mothur (1.44.3) (7) using the SILVA reference database (v132) (14) to produce count tables for phylum, class, order, family, genus, OTU, and ASV following the Schloss Lab MiSeq SOP described on the mothur website (https://mothur.org/wiki/miseq_sop/). ASV level data was also produced using DADA2 (15) to ensure consistent results with a different pipeline. Data was processed following the DADA2 pipeline, but setting pool=TRUE to infer ASVs from the whole dataset rather than per sample. The resulting ASV table was subsampled for consistency with the mothur data. The DADA2 generated ASVs performed worse than the mothur generated ASVs (DADA2 ASV mean AUROC: 0.659, p < 0.05).

### Machine Learning

Machine learning models were run with the R package mikropml (v0.0.2) (8) to predict the diagnosis category (normal vs SRN) of each sample. Data was preprocessed to normalize values (scale/center), remove values with zero or near-zero variance, and collapse collinear features using default parameters. Initially the models were run with default hyperparameters, but were expanded if the peak performance was at the edge of the hyperparameter range. Each taxonomic model taxonomic level combination (e.g. random forest on genus) was run with 100 different seeds. Each seed split the data into a training (80%) and testing (20%) set, and output performance of the training and testing as area under the receiver operating curve (AUROC).

To compare performance between taxonomic levels and models, P values were calculated as previously described (6). To compare sensitivity at 90% specificity, probabilities on the test dataset were collected for each seed and used to calculate sensitivity for specificity values ranging from 0 to 1 in 0.01 increments. The sensitivity at a specificity of 90% was pulled for each seed. The averaged ROC curves were plotted by taking the average and standard deviation of the sensitivity at each specificity value. An optional output from the mikropml package is the permuted feature importance which is quantified by iteratively permuting each feature in the model and assessing the change in model performance. Features are presumed to be important if the performance of the model, measured by the AUROC, decreases when that feature is permuted. Ranking of feature importance was determined by ordering the features based on the average change in AUROC across the 100 seeds where features with a larger decrease in AUROC are ranked higher in importance.

To quantify prevalence of the features, the number of samples with non-zero abundance was divided by the total number of samples resulting in values ranging from 0 to 1 where 0 indicates the feature is not found in any samples, 0.5 indicates the feature is found in half of the samples, and 1 indicates the feature is found in all of the samples.

All code is available at: https://github.com/SchlossLab/Armour_Resolution_XXXX_2021

## Acknowledgements

This work was supported through a grant from the NIH (R01CA215574).

## Supplemental Figures

**Supplemental Figure 1:**
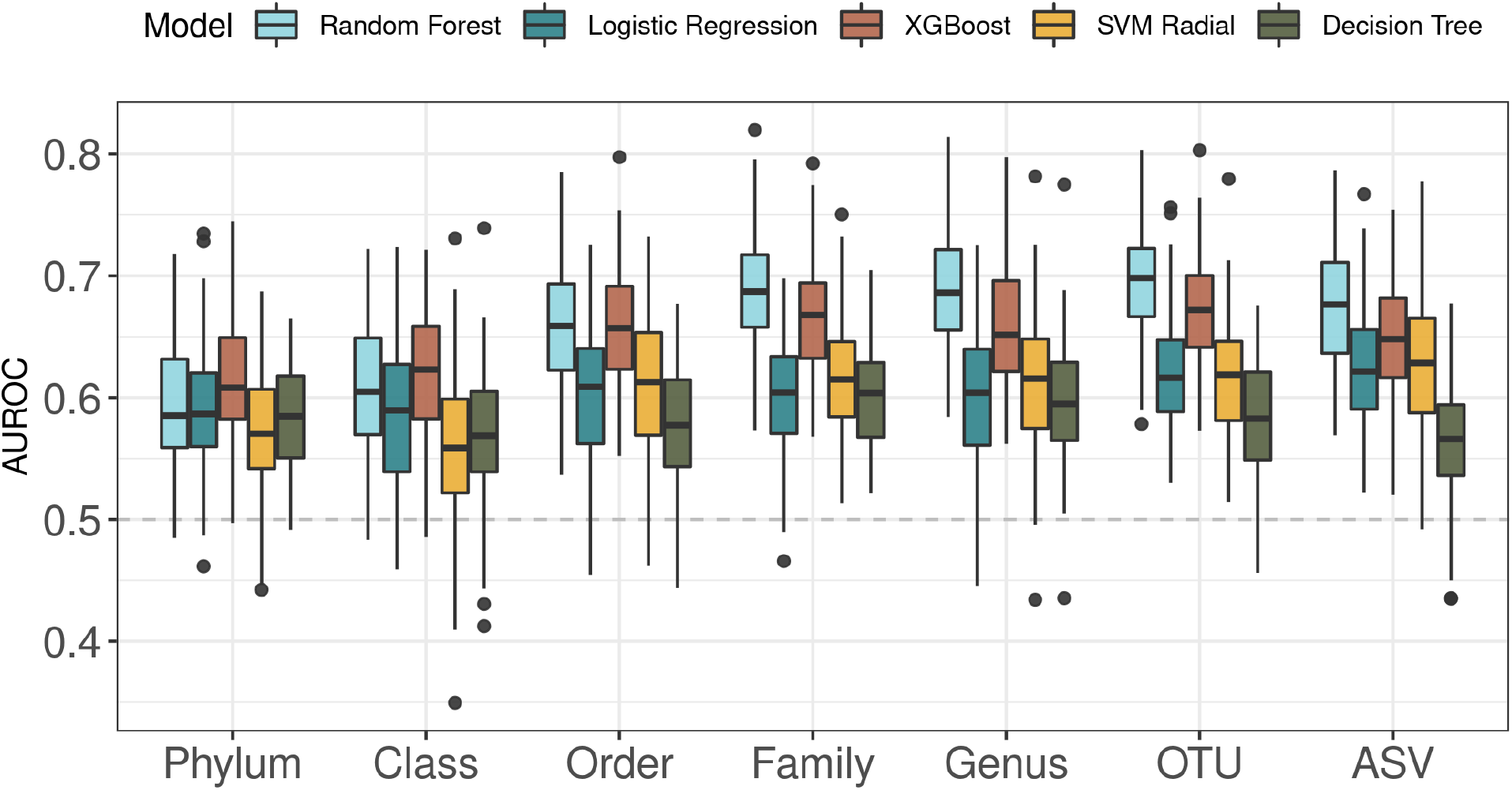
Model Performance across Taxonomy. Boxplots of AUROC values from predicting whether samples came from subjects with screen relevant neoplasias (i.e. advanced adenoma or cancer) or healthy controls across five machine learning methods including random forest, L2-regularized logistic regression (logistic regression), decision tree, gradient boosted trees (XGBoost), and support vector machine with radial basis kernel (SVM radial). Due to the random split of data into training and testing sets, each model was run across 100 seeds to account for variation in training/test datasplits.

**Supplemental Figure 2:**
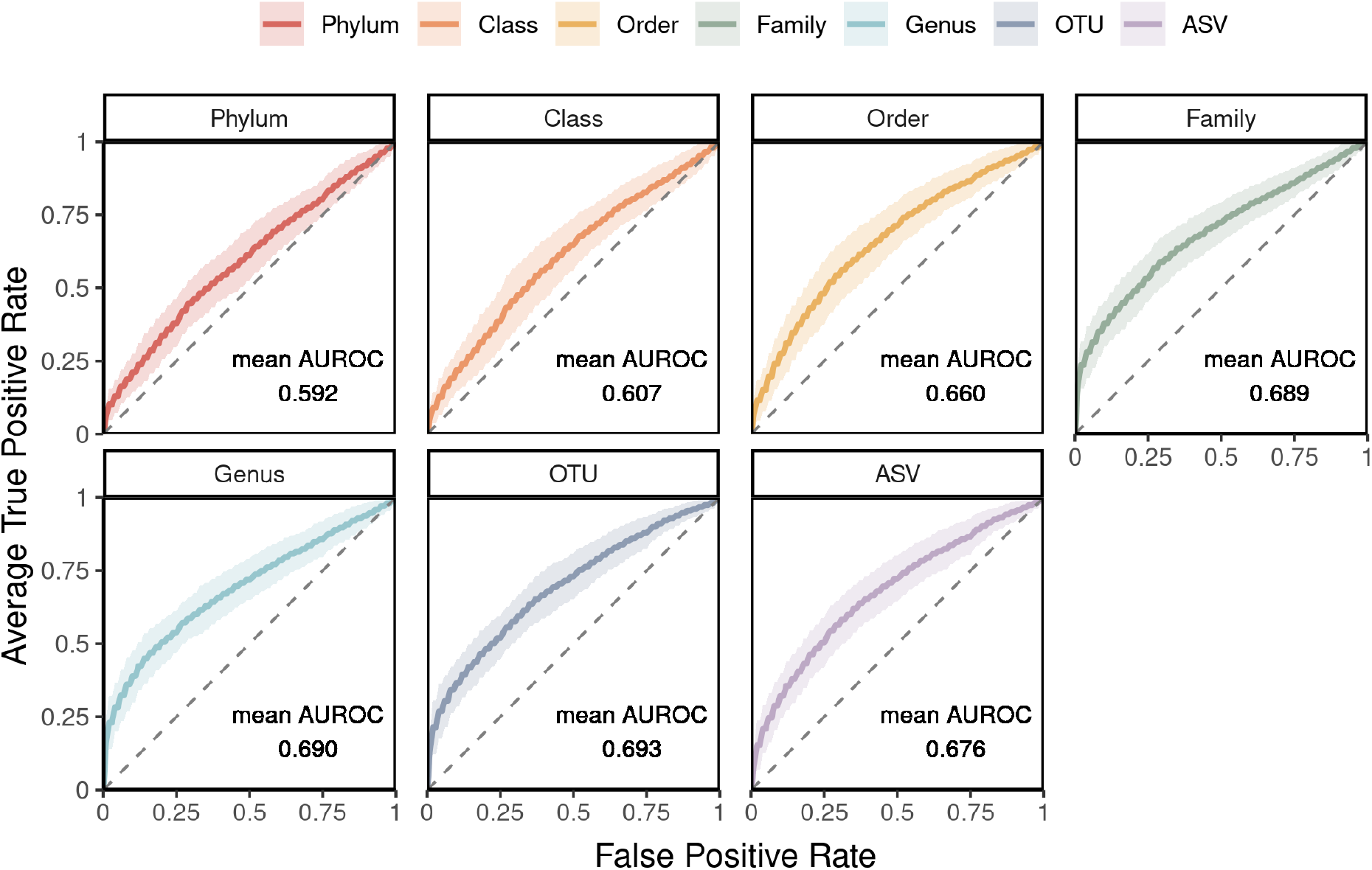
Averaged ROC curves. ROC curves with averaged true positive rate (or sensitivity) across the 100 iterations of the random forest model. The shaded region represents the standard deviation form the mean. Dashed line represents an AUROC of 0.5, which is equivalent to random classification. The mean AUROC for each taxonomic level is printed on the bottom right of the plot.

**Supplemental Figure 3:**
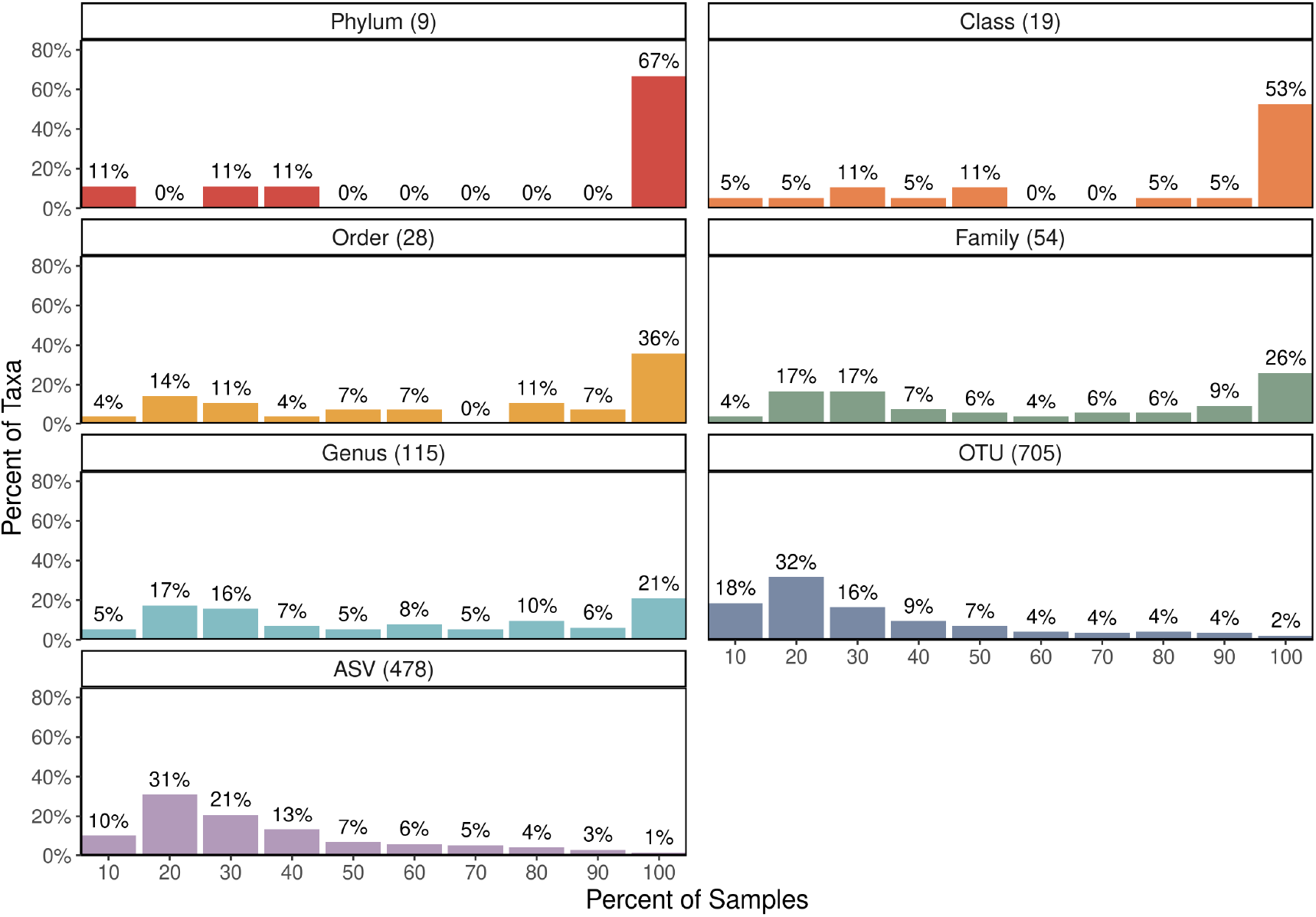
Prevalence of Taxa in Samples. Distribution of the prevalence of taxa across samples at each taxonomic level. Percent of samples is split into 10 groups where the first is for taxa present in 0 to 10% of samples, then >10% to 20% of samples, and so on. The total number of taxa for each taxonomic level after preprocessing is in parenthesis next to the title of the plot (or the name of the taxonomic level).

**Supplemental Figure 4:**
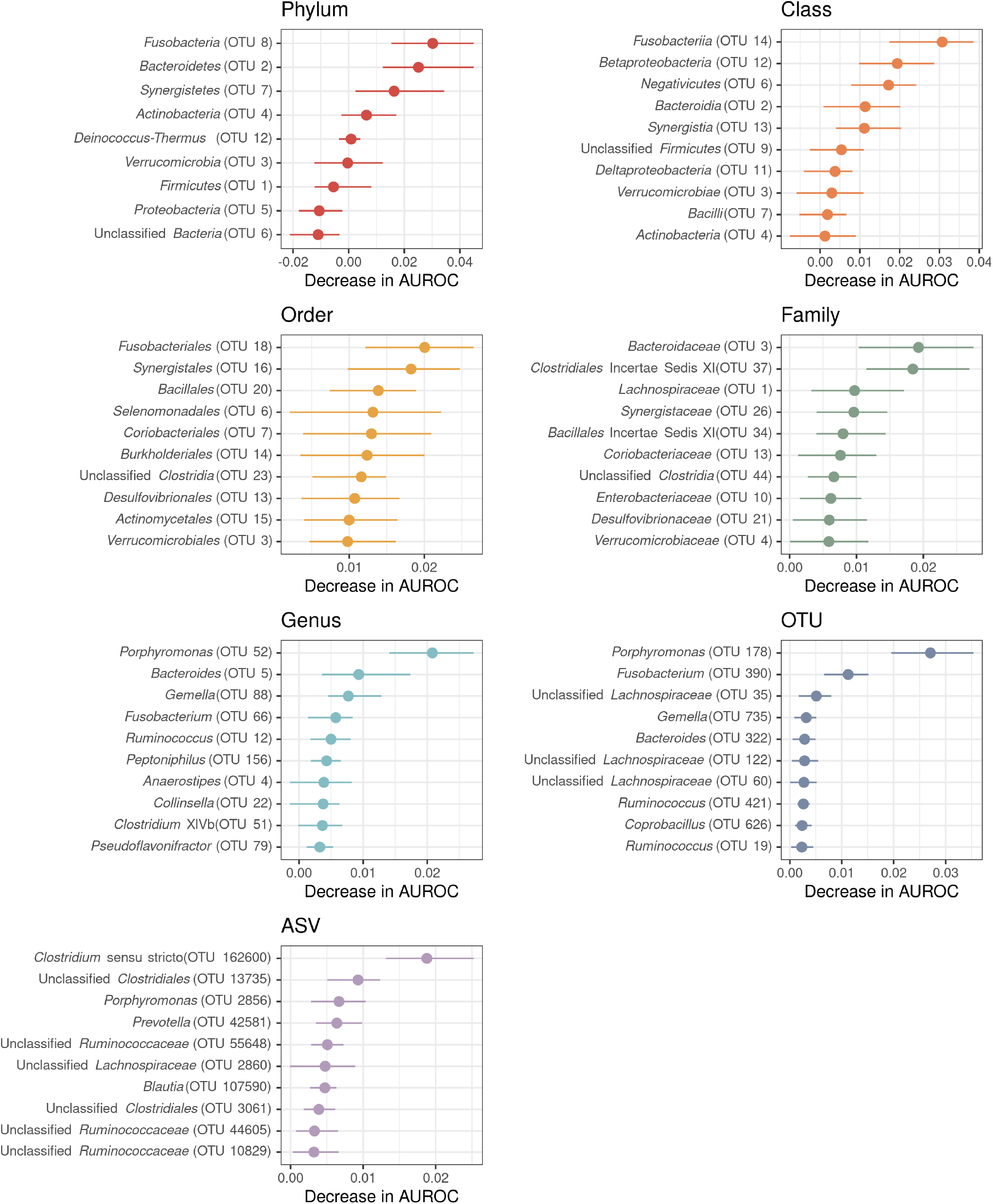
Top 10 important taxa at each taxonomic level. Summary of the 10 most important taxa for the random forest models at each taxonomic level based on the average decrease in AUROC when the feature is permuted. Dot represents the mean decrease in AUROC and the lines extending from the dot represent the standard deviation from the mean.

